# Effectiveness of FLASH vs conventional dose rate radiotherapy in a model of orthotopic, murine breast cancer

**DOI:** 10.1101/2024.12.14.628467

**Authors:** Stavros Melemenidis, Vignesh Viswanathan, Suparna Dutt, Rakesh Manjappa, Naviya Kapadia, Brianna Lau, Luis A. Soto, Ramish M. Ashraf, Banita Thakur, Adel Z. I. Mutahar, Lawrie B. Skinner, Amy S. Yu, Murat Surucu, Kerriann M. Casey, Erinn B. Rankin, Kathleen C. Horst, Edward E. Graves, Billy W. Loo, Frederick M. Dirbas

## Abstract

**Purpose:** Radiotherapy is an effective breast cancer treatment that enhances local tumor control and prolongs overall survival yet is associated with undesirable side effects which can impair quality of life. Ultra-high dose rate radiotherapy (FLASH) has been shown to induce less normal tissue toxicity while producing comparable tumor growth delay in a variety of preclinical tumor models when compared with conventional dose rate radiotherapy (CONV). However, growth delay is not a surrogate for tumor eradication, which is a critical endpoint of cancer therapy, and studies using FLASH in breast cancer are limited. We sought to evaluate whether FLASH produced comparable tumor control to CONV in a breast cancer model with tumor eradication as the primary endpoint.

**Methods and Materials:** 10^6^ cells from the radiation sensitive mammary tumor cell line Py117 were used to create non-metastatic, syngeneic, orthotopic tumors in the left 4^th^ mammary fat pad of C57BL/6J mice (n=67). Tumors were established for two distinct sequential irradiation studies (Rounds 1 and 2), utilizing either large (7.5 mm into the body) or small (5 mm) treatment tumor margins, respectively. For Round 1, mice were divided into groups with either small (20–40 mm³) or large (250–800 mm³) tumors, whereas only small tumors were included in Round 2. Tumors were irradiated with FLASH (93, 192 and 200 Gy/s) or CONV (0.08 Gy/s) using 16.6 MeV FLASH and 15.7 MeV CONV electron beams. Mice in the small tumor cohort were treated with single fractions of 20, 25, or 30 Gy. The larger tumors were treated with a single fraction of 30 Gy. Tumor eradication was determined by palpation and with histology as needed to clarify physical findings.

**Results:** Single fractions of FLASH and CONV demonstrated comparable treatment responses within matched cohorts of small and large tumors. A portion of small tumors treated with single fractions of 20 or 25 Gy were eradicated though most regrew within 2 to 3 weeks. Eradication of small tumors was best seen treated with 30 Gy and a large treatment tumor margin. These mice had no tumor regrowth at 30 days with either FLASH or CONV: however, euthanasia criteria were met at the 30-day time point due to concerns over skin toxicity for both FLASH and CONV groups. Small tumors treated with 30 Gy and a smaller treatment tumor margin had less skin toxicity with 75% of mice remaining tumor free at 48 days. 30 Gy FLASH and CONV applied to larger tumors demonstrated growth delay equally with a partial reduction in size but without tumor eradication.

**Conclusions:** FLASH and CONV produced comparable tumor control in this model of orthotopic, murine breast tumors. Single fractions of 30 Gy with both FLASH and CONV applied to small tumors achieved the highest rates of tumor eradication in particular when delivered with a wider treatment margin. Skin toxicity seen at this dose and in this location could be ameliorated with the use of multiple fractions or different tangents in future studies. Efforts at eradicating larger tumors would require testing higher single fraction doses, multiple fractions, and/or hypofractionated treatment regimens. The equivalent effectiveness between FLASH and CONV in this study of murine breast tumors supports ongoing evaluation of FLASH for use in treating human breast cancer. To this end future efforts at tumor eradication with single fraction FLASH doses with comprehensive evaluation of the toxicity of organs at risk as compared to CONV will be necessary. Additionally, studies of dose-response in a range of tumor volumes with additional breast cancer cell lines and tumors, including human xenografts, along with refined target margins, will guide future studies into the use of FLASH in the adjuvant therapy of primary human breast cancer.

## 1. INTRODUCTION

Radiotherapy (RT) is an effective treatment for invasive breast cancer (BC) and ductal carcinoma in situ (DCIS) (*1–3*). Unfortunately, there are associated side effects recognized by patients and physicians (*4–6*). Such normal tissue effects can include breast dermal and glandular fibrosis, breast shrinkage, shoulder dysfunction, higher rates of lymphedema, limited options for reirradiation, higher complication rates with implant-based breast reconstruction, and a limited ability to apply RT to larger areas such as whole liver or lung (*7–10*). In a recent study, Williams et al. noted a 47% rate of psychosocial side effects on a 15-point scale in patient reported outcomes for individuals receiving RT for breast cancer compared with 23% of patients overall treated with RT for malignancy (*5*). Radiation oncologists have ameliorated but not eliminated RT toxicity to normal tissue (NT) (*11*,*12*). RT related toxicity to NT accordingly still poses psychological dilemmas for newly diagnosed patients. Some of the most powerful advocates against RT are often previously treated patients who suffer from complications and convey their personal dissatisfaction to those newly diagnosed whom they encourage to avoid radiotherapy (*13*,*14*). Improved forms of RT for human breast cancer could increase use of breast conservation, reduce treatment related complications, improve patient reported outcomes, and further enhance tumor control and survival.

Currently, conventional dose rate radiotherapy (CONV) is delivered at dose rates less than 0.4 Gy/s (*15*). FLASH is a relatively new technique that delivers RT at ultra-high dose rates over 40 Gy/s (*16*). Evidence from preclinical models has shown that FLASH causes growth delay comparable to CONV with generally far less initial NT toxicity (*16–18*). Single fraction FLASH to the skin of the hind limb in mice at doses up to 30 and 40 Gy showed less skin toxicity than CONV (*19*). In a first in human safety and feasibility study, a single patient with cutaneous T cell lymphoma was treated successfully with FLASH (*20*) in one dose rather than the CONV hypofractionated therapy. However, a trial using single fraction FLASH to treat squamous cell carcinoma in cats was associated with late toxicity to bone in the treatment field, suggesting that FLASH may also have limitations with toxicity with longer followup (*21*). There is a paucity of preclinical FLASH studies assessing its merits against breast tumors (*22–27*). It is critical to further assess the relative benefits and risks of FLASH vs CONV to determine if FLASH should advance towards use in human breast cancer.

Multiple studies have shown that breast cancer is often associated with occult multifocal and/or multicentric disease (*28*,*29*). Radiotherapy has proven its worth in eradicating residual, occult disease after lumpectomy in several randomized clinical trials in humans and accordingly is a standard of care for most patients undergoing lumpectomy to reduce in-breast recurrences (*30–35*). In these early studies the rates of in-breast recurrence ranged from 7.5% to 14.3% with follow up extending from 5 to 20 years. More recent randomized trials of breast conservation therapy also using whole breast irradiation have demonstrated much lower in-breast recurrence rates such as 0.4% at 5.8 years in one randomized study (*36*) with another randomized trial showing a recurrence rate of 3.9% at 10 years (*37*). Decreased in-breast recurrence rates are likely a result of several factors, including improved preoperative imaging such as breast MRI, improved systemic therapy, and advances in radiation treatment planning (*1*). With respect to breast MRI, considered the most sensitive test for identifying occult breast cancer (*38*), we found that MRI was accurate down the level of lesions 4 mm in size: more findings under 4 mm were associated with false positive findings (*39*). This suggested to us that 3 mm was a reasonable baseline to use in assessing the effectiveness of radiotherapy for eradication of subclinical, occult breast cancer remaining after lumpectomy in the current era of breast conservation where MRI is commonly used prior to breast conservation surgery/lumpectomy.

Initial experience with FLASH *vs* CONV in breast cancer is promising. A preclinical study using a xenograft model with HBCx-12a BC tumors, an aggressive TNBC phenotype (*16*), demonstrated delay in tumor growth, but not eradication. Growth delay, akin to a partial clinical response after neoadjuvant chemotherapy, is not a clear surrogate for eradication. Eradication is a necessary oncologic endpoint for successful radiotherapy. The most common assessment of comparative tumor eradication between different radiotherapy techniques is tumor control dose 50 (TCD_50_), which refers to the radiation dose required to achieve local control of a tumor in 50% of the cases. Such studies require a large number of mice. The only study to date evaluating TCD_50_ was performed in a heterotopic, subcutaneous (hind limb), mammary carcinoma mouse model with tumor volumes of 200 mm^3^ using proton pencil beam and showed comparable TCD_50_ of around 50 Gy for both FLASH and CONV (*40*). To further advance knowledge of the ability of FLASH to control tumor growth in a more clinically relevant scenario, there is rationale for testing FLASH using complete eradication of all tumors as an endpoint and using an orthotopic location.

A separate study using a murine model of glioblastoma demonstrated that tumor eradication with FLASH led to long term immunity to subsequent tumor challenge (*41*). This further suggests merit to identifying techniques for orthotopic, single fraction tumor eradication as might be used after lumpectomy as a potential opportunity for induction of long-term immunity.

For this pilot study, we focused our efforts on eradication of small tumors 20–40 mm^3^ but also treated larger tumors 250-800 mm^3^ which is more in keeping with results from other preclinical studies. We hypothesized that if FLASH were as effective as CONV in eradicating small tumor nodules, approximating residual occult disease after lumpectomy in human breast cancer, this would provide significant justification for further evaluation of FLASH towards use in human breast cancer. We also hypothesized that FLASH would be as effective as CONV in producing growth delay associated with a temporary reduction in tumors size with larger tumors to further confirm the effectiveness of FLASH *vs* CONV seen by other investigators in preclinical breast cancer tumor models.

## 2. METHODS AND MATERIALS

### 2.1. Animals

All animal experiments and procedures were approved by the Institutional Animal Care and Use Committee of Stanford University in accordance with institutional and NIH guidelines. Six- to eight-week-old female C57BL/6 mice were obtained from The Jackson Laboratory (Bar Harbor, ME). Standard animal care and housing were provided by the Stanford University School of Medicine and is under the care and supervision of the Department of Comparative Medicine’s Veterinary Service Center. Mice were maintained on the irradiated Envigo Teklad diet containing 18% protein and 6% fat.

### 2.2. Orthotopic mouse model

A radiation sensitive mammary tumor cell line, Py117, derived from the transgenic model of the mouse mammary tumor virus promoter driving the polyoma middle T antigen (MMTV-PyMT), was chosen as the tumor model. Py117 efficiently forms non-metastatic orthotopic tumors in C57BL/6 mice (*42*). Py117 cells were cultured using F12K media containing 5% fetal clone II (Hyclone), MITO (1:1,000 dilution, BD Biosciences), 50 mg/mL gentamicin and 2.5 mg/mL amphotericin B (*42*). Injections were prepared with 10^6^ cells suspended in 50 µL sterile PBS. Ten- to eleven-week-old healthy C57BL/6 female mice (n = 67) were inoculated in the left 4^th^ mammary fat pad while under anesthesia (induction 3% and maintenance 1.5% isoflurane in pure O_2_). Tumor inoculations were performed in two sequential distinct experiments, Round 1 with 7.5 mm treatment margin and Round 2 with 5 mm treatment margin. In Round 1, mice were divided into groups with either small tumor sizes (20–40mm^3^; n = 16) or large tumor sizes (250-800 mm^3^; n = 12), while in Round 2 only small size tumors were irradiated (20–40mm^3^; n = 32). For mice in the small tumor cohorts, tumor nodules were allowed to grow for a week, while the larger tumors were allowed to grow for 4 weeks. A detailed experimental design for both rounds is outlined in **Table 1**.

**Table 1.** Experimental Irradiated Groups of Breast Cancer Orthotopic Tumor-Bearing Mice, Irradiated with Either FLASH or CONV Dose Rates in Two Irradiation Rounds with Different Planning Treatment Volumes.

| Round 1 |  |  |  |  |  |  |  |
| --- | --- | --- | --- | --- | --- | --- | --- |
| Mode | Tumor Volume<br>[mm<sup>3</sup>] | Dose<br>[Gy] | No. mice | Collimation<br>[cm<sup>2</sup>] | PTV<sup>a</sup><br>[cm<sup>2</sup>] | SSD<sup>b</sup><br>[cm] | Missed<br>Pulses* |
| FLASH | 20-40 | 20 | 4 | 2 × 2 | 0.75 × 2.00 | 18.7 | 1 |
| FLASH | 20-40 | 30 | 4 | 2 × 2 | 0.75 × 2.00 | 18.7 | 0 |
| FLASH | 250-800 | 30 | 6 | 2 × 2 | 0.75 × 2.00 | 18.7 | 0 |
| CONV | 20-40 | 20 | 4 | 2 × 2 | 0.75 × 2.00 | 147.5 | - |
| CONV | 20-40 | 30 | 4 | 2 × 2 | 0.75 × 2.00 | 147.5 | - |
| CONV | 250-800 | 30 | 6 | 2 × 2 | 0.75 × 2.00 | 147.5 | - |
| Round 2 |  |  |  |  |  |  |  |
| Mode | Tumor Volume<br>[mm<sup>3</sup>] | Dose<br>[Gy] | No. mice | Collimation<br>[cm<sup>2</sup>] | PTV<sup>a</sup><br>[cm<sup>2</sup>] | SSD<sup>b</sup><br>[cm] | Missed<br>Pulses* |
| FLASH | 20-40 | 20 | 4 | 2 × 2 | 0.50 × 2.00 | 18.7 | 0 |
| FLASH | 20-40 | 25 | 8 | 2 × 2 | 0.50 × 2.00 | 18.7 | 1 |
| FLASH | 20-40 | 30 | 4 | 2 × 2 | 0.50 × 2.00 | 18.7 | 0 |
| CONV | 20-40 | 20 | 4 | 2 × 2 | 0.50 × 2.00 | 18.7 | - |
| CONV | 20-40 | 25 | 8 | 2 × 2 | 0.50 × 2.00 | 18.7 | - |
| CONV | 20-40 | 30 | 4 | 2 × 2 | 0.50 × 2.00 | 18.7 | - |

<sup>a</sup> Planning treatment volume; margin area within the mouse positioner frame (abdomen exposure).
<sup>b</sup> Source to surface distance.
\* Animals irradiated with missed pulses were excluded from the analysis.

### 2.3. Tumor volume measurements

Tumor measurements were acquired without anesthesia. Mice were scruffed and tumor measurements acquired using calipers. Tumor volumes were derived from two orthogonal measurements using the ellipsoid approximation formula (volume = 0.5 × length × width^2^). Tumor measurements were initiated when tumors became palpable, and thereafter, three days per week.

### 2.4. Dose target groups

In Round 1, small tumor cohorts were treated with single fractions of 20 or 30 Gy target dose of FLASH or CONV (n = 4 per group). The large tumor cohort was treated with a single fraction of 30 Gy target dose of FLASH or CONV (n = 6 per group). Seven control mice were left untreated and used to confirm tumor growth in the absence of irradiation. In Round 2, only small size tumors were treated (n = 36). The small size groups treated in Round 1 with 20 or 30 Gy target dose with either FLASH or CONV were also repeated in Round 2 (n = 4 per group). Additionally in Round 2, a target dose group at dose midpoint between 20 and 30 Gy was treated with 25 Gy target dose of FLASH or CONV (n = 8 per group). A detailed experimental design for both rounds is outlined in **Table 1**.

### 2.5. Stereotactic mouse positioner and beam collimator

During positioning and irradiation, mice were anesthetized with a mixture of ketamine (100 mg/kg) and xylazine (10 mg/kg) injected into the peritoneum. Immediately after irradiation, mice were placed on a warming blanket until they recovered from anesthesia. They were then returned to their standard housing environment. All 3D-prints were designed in Fusion 360^®^ (Autodesk, San Rafael, CA, USA), 3D computer-aided design (CAD) files were edited with Ultimaker Cura v.4.3.1 and printed with Ultimaker S5^®^ (New York, NY, USA) using polyactic acid (PLA).

The mice were placed in a 3D-printed stereotactic mouse positioner with a 1.5 x 2.0 cm lateral opening. The mice were immobilized, allowing part of the abdomen to protruding outside the wall of the positioner (**Fig. 1A**). The mice tumors were centered at the lower part of the lateral opening of the mouse stereotactic positioner and the lower bodies of the mice were gently pushed towards the opening of the positioner using paper tissue. The stereotactic mouse positioner is placed on the top of the beam collimator with a 2.0 x 2.0 cm radiation exposure field (**Fig. 1B**). The stereotactic mouse positioner interlocks on the top of the collimator, which provides serial 2.5 mm lateral slots across the field of irradiation field. This setting allows for irradiation exposures at different depths within the stereotactic positioner, providing an option for various treatment tumor margins (**Fig. 1C**). The beam collimator is designed for electron irradiation geometry with floor-to-ceiling beam orientation, with a 3-cm-thick layer of aluminum oxide powder (Al_2_O_3_; 99.99% trace metals basis; Sigma-Aldrich, St Luis, MO, USA) for electron attenuation, in tandem with a 1-cm-thick layer of tungsten spheres (2 mm diameter; TPW, WY, USA) for reduction of Bremsstrahlung radiation produced from the collimator’s material (0.2% Bremsstrahlung radiation leakage; **Fig. 1D**).

**FIG 1.**
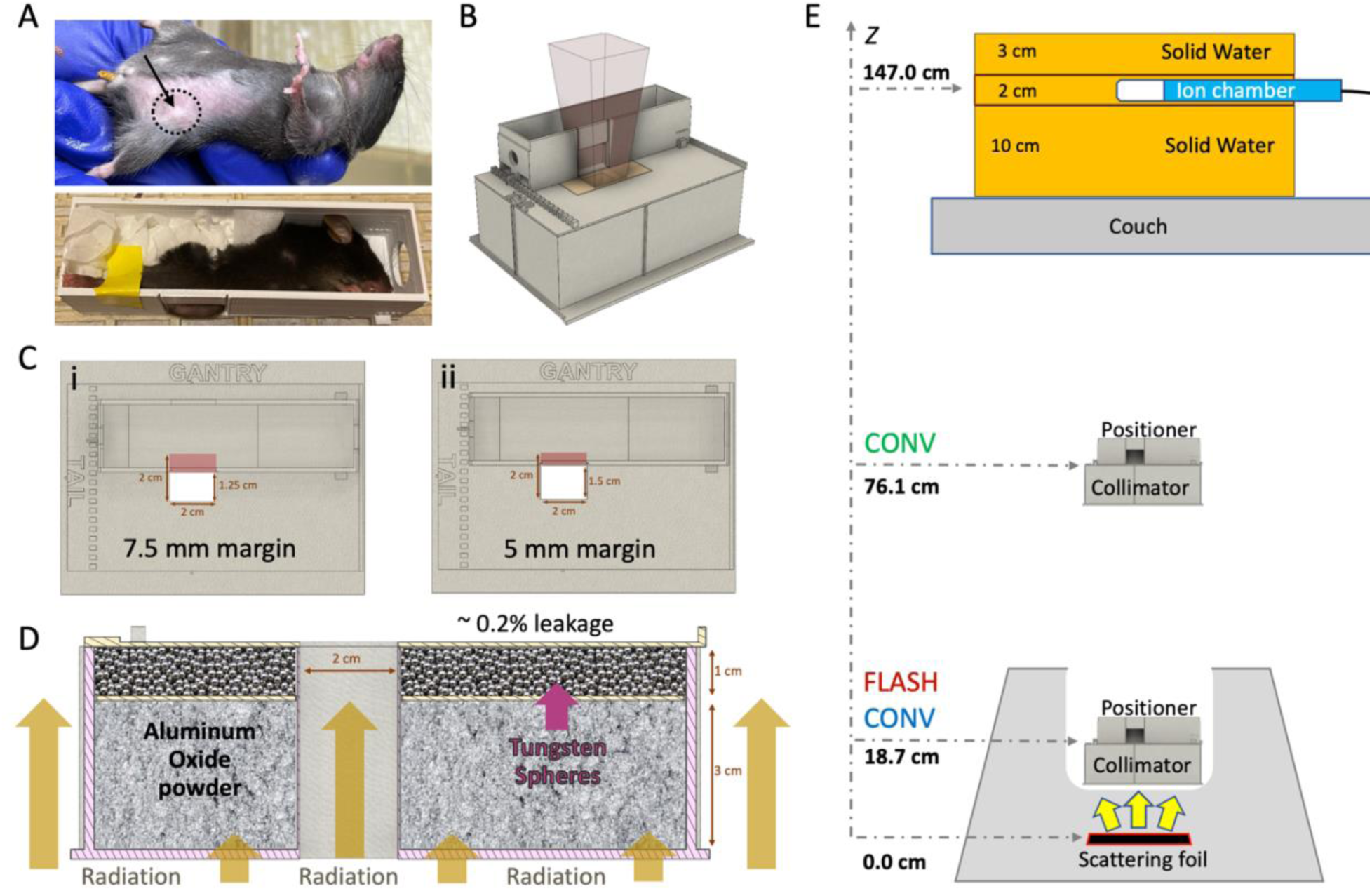
*In vivo* breast cancer orthotopic tumor irradiations with FLASH or CONV dose rates. (**A**) Breast cancer orthotopic tumor-bearing mouse with approximately 30 mm^3^ tumor at the 4^th^ mammary fat pad (top; black dashed circle and arrow). An anesthetized mouse placed inside the mouse positioner frame with the tumor centered at the bottom of the lateral opening and immobilized with paper tissue (bottom). (**B**) 3D computer-aided design (CAD) files illustrating the positioning of the mouse positioner on the collimator, the positioning of the radiochromic film during irradiation. (**C**) Top view of the collimator featuring the mouse positioner in two treatment tumor margins: (**i**) Round 1, positioned to expose a 7.5 × 0.2 mm² area within the abdominal tissue or (**ii**) Round 2, positioned to expose a 5.0 × 0.2 mm² area within the abdominal tissue. (**D**) 3D CAD file of the 2.0 × 2.0 cm^2^ collimator presented with a lateral cross-section, illustrating mouse radiation shielding. The collimator is filled with a 3-cm-layer of aluminum oxide to stop the electrons and a 1-cm-layer of tungsten spheres (2.0 mm diameter) to absorb Bremsstrahlung radiation and efficiently shield the rest of the animal’s body (Bremsstrahlung radiation leakage ∼0.2%). (**E**) FLASH and CONV beam geometries. For FLASH irradiations, the collimator and mouse positioner frame are placed inside the treatment head and the beam entrance surface of the mouse is 18.7 cm from the scattering foil. For CONV irradiations, at Round 1, the beam entrance surface of the mouse was 76.1 cm (unmatched geometries) from the scattering foil, and at Round 2 at 18.7 cm (matching geometries with FLASH). The pulses delivered are monitored using an ion chamber, measuring the Bremsstrahlung tail at 11.0 cm solid water depth, and is located 147.5 cm from the scattering foil.

### 2.6. Irradiation study design

Tumor-bearing mice (n = 60) were irradiated with either FLASH (94, 193 or 200 Gy/s) or CONV (0.136 or 0.146 Gy/s) dose rates using approximately 16 MeV (E_0_ = 16.6 MeV FLASH and 15.7 MeV CONV; **Table 2**) electron beams from a previously described configuration of a clinical Varian Trilogy (*43*,*44*) and a microcontroller-based (Red Pitaya, Slovenia, Europe) pulse control methodology (*43*). Radiotherapy treatments were administered in two sequential distinct experiments, Round 1 and 2, with some key different between them: a) in Round 1 the source-to-surface distance (SSD; scattering foil to mouse surface) for FLASH beam geometry was 18.7 cm compared to 76.1 cm SSD for the CONV beam geometry (**Fig 1E**, red and green notation), as described previously (*45*,*46*). On Round 2, due to improvements in our configuration the beam geometry mismatch was resolved and all experiments thereafter implemented same geometry between modalities. (**Fig 1E**, red and blue notation) (*47*). b) In Round 1, the mouse stereotactic positioner was set to a large treatment tumor margin with 7.5 mm radiation exposure within the abdomen (**Fig. 1C.i**) to compensate for the geometric disparity. In Round 2, the geometric disparity was resolved and due to concerns of radiation toxicity the treatment tumor margin was reduced to 5.0 mm exposure within the abdomen (Round 2; **Fig 1C.ii**). c) In Round 1, only even number of Gy were used in the target dose groups (20 and 30 Gy) and therefore 2 Gy pulses were used, while in Round 2, a dose midpoint 25 Gy dose target group was introduced that required implementation of 1 Gy pulses. **Table 1** shows all experimental groups and their associated beam geometry parameters and **Table 2** outline the beam characteristics and the pulse delivery design for each modality.

**Table 2.** Beam Parameters for Experimental Beam Delivery with FLASH or CONV Dose Rates.

| Mode | FLASH | CONV |
| --- | --- | --- |
| **Beam energy [MeV]** | 16.60 | 15.73 |
| **Target dose [Gy]** | 20, 25, 30 | 20, 25, 30 |
| **Delivered pulses** | 10, 15, 30 | 1901, 4763, 9526, 19051 |
| **Pulse rate [Hz]** | 90 | 72, 90 |
| **Dose per pulse [Gy]** | 1, 2 | 1.5E-3, 2.00E-3 |
| **Dose rate [Gy/s]** | 93.75, 192.86, 200 | 0.08 |
| **Pulse length [s]** | 3.75E-6 | 3.75E-6 |
| **Intra-pulse dose rate [Gy/s]** | 2.67E+5 | 400, 533 |

### 2.7. Dosimetry

Absolute doses were determined at the surface of the beam collimator using radiochromic film (EBT3 Galfchromic, Ashland Inc., Wayne, NJ; **Fig. 2A,B**) and exit charge measurements at the Bremsstrahlung tail of the electron beam using an ion chamber (Farmer^®^ Chamber, PTW Model TN30013, Boonton, NJ; **Fig. 1E**). During each mouse irradiation, one 2.4 x 5.1 cm piece of radiochromic film per mouse was placed under the stereotactic mouse positioner, on the top of the beam collimator, centered on the 2.0 x 2.0 cm irradiation field (**Fig. 2C**). Films were allowed to self-develop for more than 24 hours post irradiation before scanned at 72 dots per inch resolution. An average optical density (OD) of the irradiated area was converted to absorbed dose using a predetermined relationship of OD *vs.* dose gradient according to the recommendations of the manufacturer (*48*). Subsequently, using the associated exit charge reading from the ion chamber for each mouse, an average of nC/Gy was determined for each group. The absolute dose per mouse was then calculated using the exit charge reading, which reduced the inherent variation of the film readings (< 0.5 % variation of ion chamber readings *vs.* < 3 % from radiochromic film). The beam profiles of the collimation were assessed at the surface of the collimator in X and Y direction for both FLASH and CONV geometries using experimental films (**Fig. 1E**; **Fig. 2B–E**). The percentage dose depth (PDD) distributions of the beam were assessed for both FLASH and CONV geometries using sagittal films, oriented parallel to the beam (Z direction), sandwiched between solid water (**Fig. 2F**).

**FIG 2.**
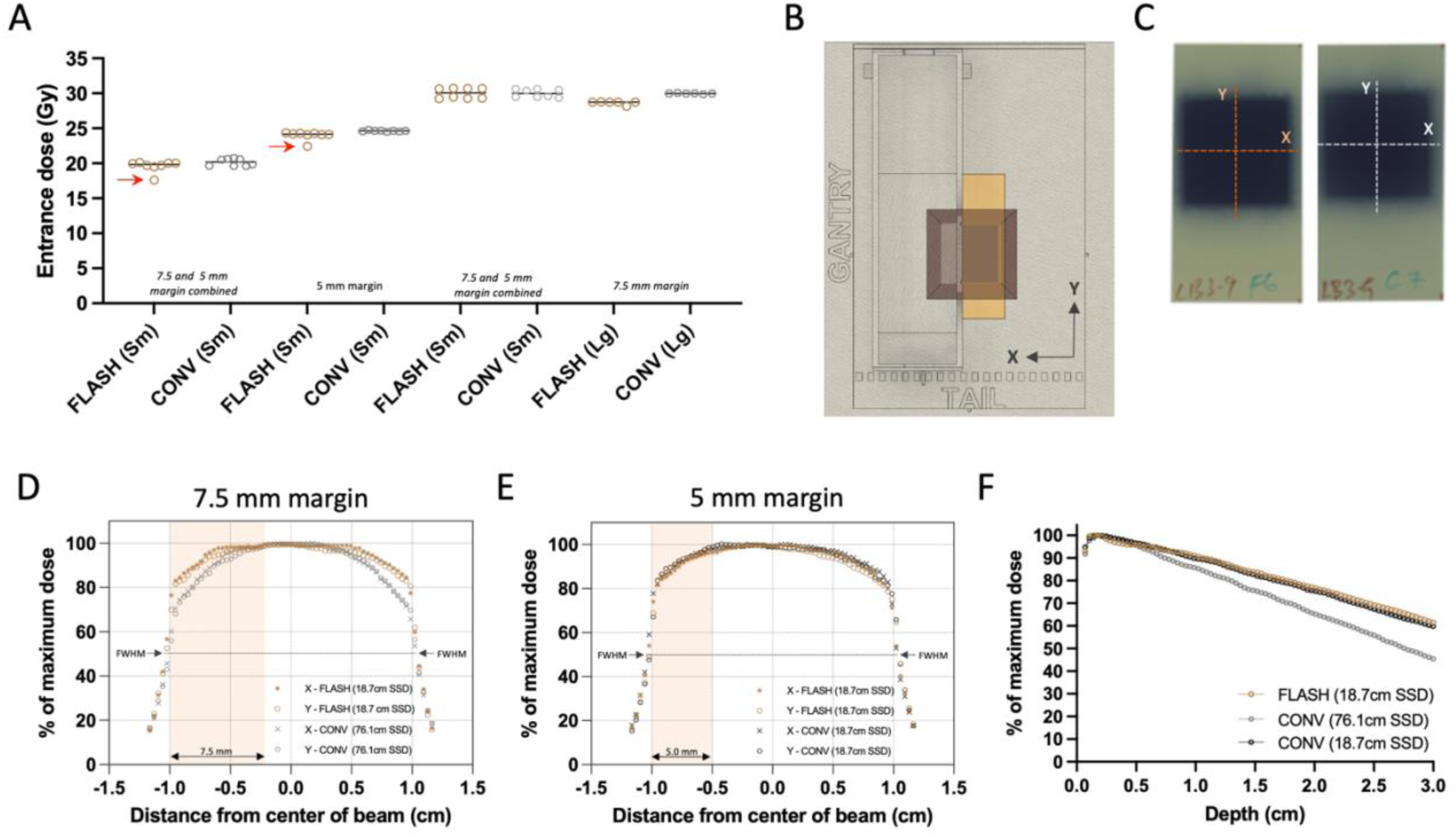
Radiochromic film dosimetry of delivered doses and film-derived collimator characterization. (**A**) Grouped scatter plot of film-derived mean dose of all animals combined (Sm for small 20–40 mm³ and Lg for large 250–800 mm³ tumor volumes). Red arrows indicate missed pulses, which resulted in elimination of these animals from the analysis. (**B**) Top view of experimental set up illustration the film positioning and exposure. (**C**) Representative exposed films with either FLASH or CONV 25 Gy single fraction target dose group, illustrating the direction of the beam profiles. (**D**) Film-derived X (transverse) and Y (craniocaudal) profiles of the unmatched geometries between FLASH and CONV with 7.5 mm treatment margin from Round 1 of irradiations, and (**E**) profiles from the matched geometries with 5 mm treatment margin from Round 2. Salmon highlights represent the margin of exposure in X direction. (**F**) Film-derived percentage depth dose curve (PDD) from FLASH and both geometries of CONV configurations, using films parallel to the direction of the irradiation beam. Overall, the delivered doses between FLASH and CONV were comparable and despite the difference in source-to-surface (SSD), profiles and PDDs of the two modalities remain comparable within the tumor volumes.

### 2.8. Beam homogeneity and dose distribution

The examination of the experimental films’ lateral profiles revealed a symmetrical distribution in the X and Y axes for both FLASH and CONV geometries, as depicted in **Fig. 2D**. In the first round of irradiations, the initial setup featuring varying beam geometries—70.2 cm for FLASH and 19.2 cm for CONV surface distance from the scattering foil, as shown in **Fig. 1E**—the full width half maximum (FWHM) of the profiles corresponded closely with the predefined field dimensions: 4.2% and 3.8% for FLASH, and 2.3% and 2.1% for CONV in the X and Y directions respectively (**Fig. 1E** and **Fig. 2C**). A notable 14.5% difference in dose profiles at the field’s perimeter prompted the incorporation of a wide treatment margin of 7.5 mm within the mouse stereotactic positioner, yielding a minimal 0.9% dose variance at the positioner’s lateral aperture, where the tumor protrudes (**Table 1**; **Fig. 1C.i** and **Fig. 2D**).

During the second round of irradiation, beam configurations were harmonized for both FLASH and CONV to a 19.2 cm surface distance from the scattering foil (**Fig. 1E**). Here, the FWHM values measured 3.2% and 3.5% for FLASH, and 3.5% and 3.4% for CONV along the X and Y axes respectively, aligning with the stipulated field size (**Fig. 1E** and **Fig. 2E**). At this stage, the dose discrepancy at the radiation field’s edge was reduced to 4.2%. Consequently, a 5 mm treatment tumor margin was established, which exhibited a 1.7% difference in dose between the FLASH and CONV modalities at the lateral entry point of the positioner where the tumor emerges (**Fig. 1C.ii** and **Fig. 2E**).

### 2.9. Radiation doses

**Figure 2A** illustrates the film-derived, charge-weighted absorbed doses on the surface of the radiation shield. There was slight dose variance within the groups irradiated separately in Round 1 and 2 (with 4 mice per round undergoing 20 and 30 Gy target dose treatments on smaller tumors) when considering FLASH or CONV treatments individually. However, the overall dose levels between the FLASH and CONV groups were in alignment. When averaging the measured doses from both rounds for small tumors (this includes the 20 Gy and 30 Gy cohorts that were irradiated in two rounds, as well as the 25 Gy cohort that was irradiated in one round), the absorbed doses were closely matched between FLASH (19.9 ± 0.24, 24.0 ± 0.11, and 30.0 ± 0.73 Gy) and CONV (20.2 ± 0.51, 24.6 ± 0.08, and 30.0 ± 0.58 Gy) modalities, with a difference of less than 3% noted. A more pronounced variation of 4.5% was observed in the large tumor-bearing mice targeted with 30 Gy dose, where the FLASH group had an average absorbed dose of 28.6 ± 0.25 Gy in comparison to the CONV group’s 30.0 ± 0.06 Gy. In the target dose groups of 20 and 25 Gy, one specimen from each group received one fewer 2 Gy pulse than prescribed, as can be discerned in Fig. 2A (indicated by red arrows). Mice receiving a lower number of pulses were omitted from subsequent analysis reported below.

## 3. RESULTS

### 3.1. FLASH shows comparable tumor control to CONV against small tumors at 20, 25, and 30 Gy, with 30 Gy eradicating most small tumors

Smaller tumors, averaging 30 mm³ and treated with a single 20 or 25 Gy dose, exhibited similar levels of tumor regression with FLASH and CONV with tumor eradication in some and tumor regrowth in others. From the total of 30 tumors (n=7 per group for 20 Gy and 25 Gy FLASH, n=8 per group for 20 Gy and 25 Gy CONV), 26 tumors initially responded with complete remission (unmeasurable) while one tumor per group remained palpable. Overall, tumors which were not eradicated regrew to < 8 mm^3^ within 21 to 28 days of irradiation (Fig. 3A and 3B). For the 20 Gy irradiation groups, 29% of tumors receiving FLASH and 43 % tumors receiving CONV group (1 death) remained in complete remission on day 46. The remainder grew past their original volume, with one exception from each group that had half the original volume (Fig. 3A). At the same time point, for the 25 Gy FLASH group, 83% of tumors (1 death) regrew to their original volume, with exception of 1 tumor that had 60% of its original volume, while 50% (2 deaths) for the 25 Gy CONV group regrew above their original volume, with exception of 2 tumors that had < 50% of its original volume (Fig. 3B). There was no significant difference in the growth curves between 20 and 25 Gy for either radiotherapy modality group by day 46, as depicted in Fig. 3A**,B**. Of note, the tumors treated at 20 Gy had a wider treatment margin of 7.5 mm *vs* tumors treated at 25 Gy which had only a 5.0 mm treatment margin. There was no clear cause of death in the mice that died during the study period as they exhibited neither skin toxicity nor signs of gastrointestinal toxicity such as diarrhea.

**FIG 3.**
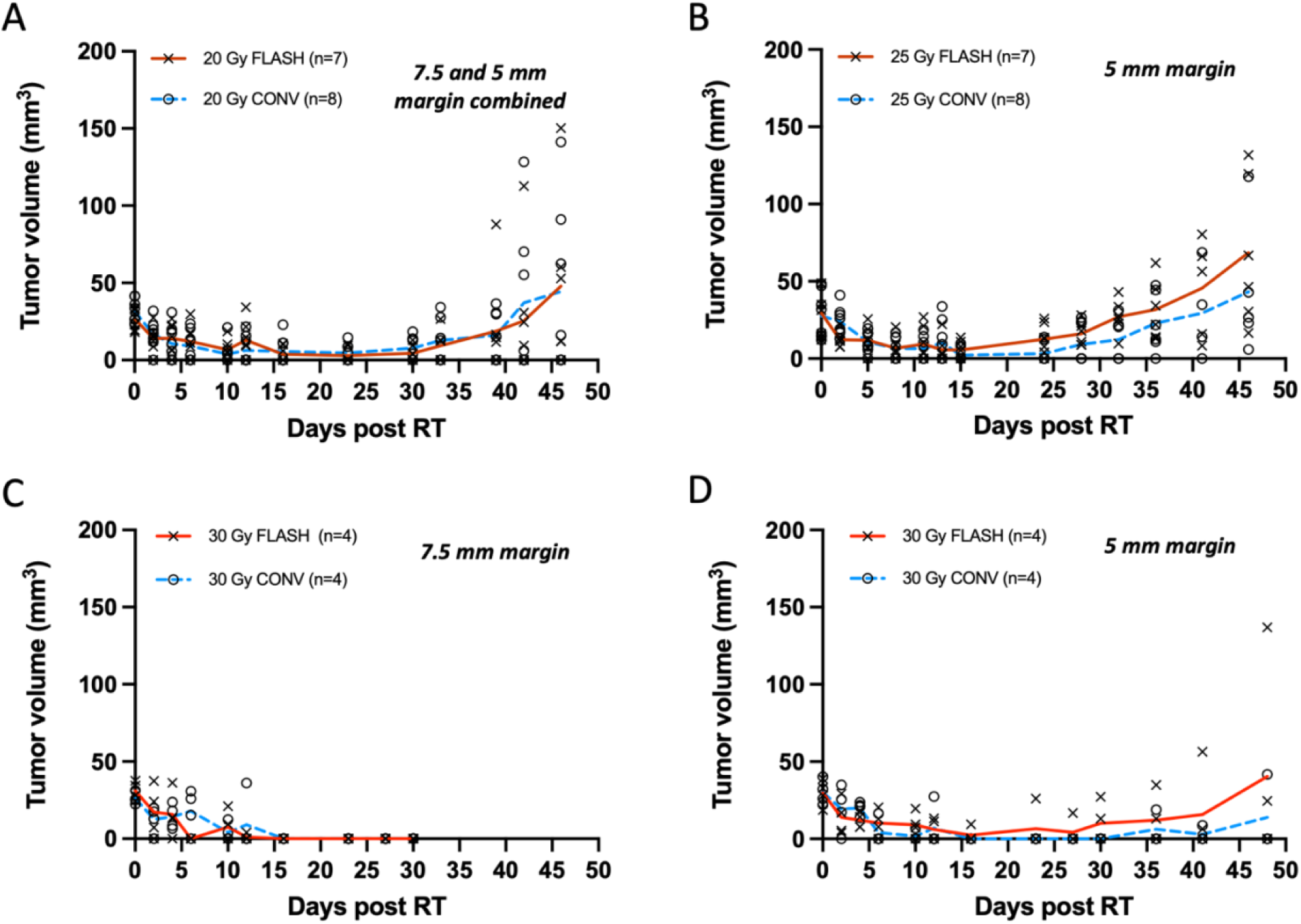
(**A-D**) Tumor measurements with calipers plotted as response curves of breast cancer orthotopic tumors of small volume (20-40 mm^3^) irradiated with 20, 25 and 30 Gy single fraction with either FLASH or CONV dose rates. (**A**) Tumor volumes from animals irradiated with 7.5 and 5 mm treatment margin combined (n=7 FLASH, n=8 CONV). FLASH group had one animal excluded due to a missed pulse. Tumors were controlled for the first 4 weeks and regressed thereafter, without significant statistical difference between FLASH and CONV groups. (**B**) Tumors targeted with 25 Gy and 5 mm treatment margin (n=7 FLASH, n=8 CONV). FLASH group had one animal excluded due to a missed pulse. Tumors were controlled for the first 4 weeks and regressed thereafter, without significant statistical difference between FLASH and CONV groups. (**C**) Tumor targeted with 30 Gy and 7.5 mm treatment margin (n=4 per group) were controlled by day 30 but due to severe tissue toxicity the experiment was terminated. (**D**) Tumors targeted with 30 Gy and 5 mm treatment margin were controlled for the first 4 weeks and by day 48 only one animal per group had measurable tumor regrowth. Overall, there was no significant difference between FLASH and CONV in tumor growth delay or eradication of small tumor volumes (20–40 mm³) with single fractions of 20, 25 and 30 Gy.

Treatments with a 30 Gy single dose, similar to the 20 Gy group, were also carried out in two rounds, as presented in Fig. 3C and 3D. In Round 1, the disparate beam geometries for FLASH and CONV required a 7.5 mm margin to account for the difference at the field edge (**Table 1**; Fig. 1D**.i**). Both modality groups (n = 4 per group) exhibited complete remission by two weeks, which persisted up to four weeks. Nonetheless, due to skin radiotoxicity, the study concluded at day 30 as euthanasia criteria were met and all mice in the cohort were sacrificed due to ulceration or toxicity concerns. (Fig. 3C). In Round 2, with matched beam geometry to minimize radiotoxicity, a smaller 5 mm margin was employed (**Table 1**; Fig. 1D**.ii**). Following the 30 Gy treatment with a 5 mm margin, both modality groups (n = 4 per group) exhibited complete remission by two weeks. By day 48, 50 % of tumors from the FLASH group and 66 % of tumors (1 death) from the CONV group remained in complete remission. At the same time point, one tumor from the FLASH group had regrown past its original volume, and one had regrown to 75% of its original volume, while in the CONV group, one mouse’s tumor had regrown to its original size (Fig. 3D). Histological verification indicated that one nodule in the FLASH group was not a tumor as suspected (**Supplementary Fig. 1A**) but rather an enlarged lymph node, as shown in **Supplementary Fig. 1B**. Regardless of the histological adjustments, there was no significant difference in tumor growth between the 30 Gy FLASH and CONV groups.

Overall, of the 20, 25, and 30 Gy doses, the 30 Gy dose led to greater tumor eradication, and a 7.5 mm treatment margin was associated with better eradication than the 5 mm treatment margin.

### 3.2. FLASH is as effective as CONV in delaying growth of larger tumors

In the initial week, unirradiated Py117 orthotopic tumors typically reach a volume of about 30 mm³, escalating to ∼ 635 mm³ by the fourth week, and surpassing 1000 mm³ in the fifth week, as shown in Fig. 4A. While smaller tumors display a consistent size range, larger ones exhibit greater variability in size. Tumors of larger size, when treated with a 30 Gy dose with either FLASH or CONV radiation, demonstrated similar regression within the first two weeks post-irradiation, followed by a parallel pattern of regrowth. Despite the FLASH group receiving a slightly reduced dose of 28.6 Gy compared to the 30 Gy for the CONV group, tumor control was equivalent between the two modalities. The study concluded after three weeks since the criteria for euthanasia were met due to ulcerations associated with large tumors and/or skin toxicity, which included alopecia, affecting three mice in the FLASH group and four in the CONV group. It was determined that a 30 Gy irradiation dose was insufficient for the complete eradication of larger tumors, as depicted in Fig. 4B.

**FIG 4.**
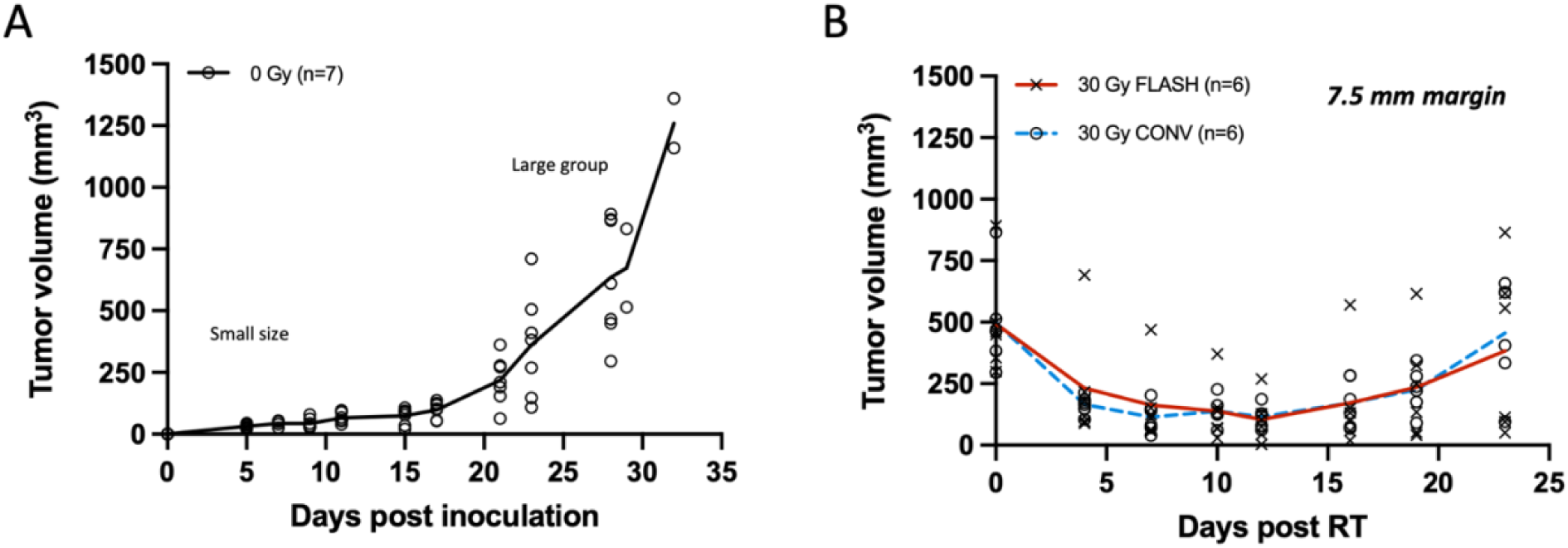
(**A**) Tumor growth curve of unirradiated breast cancer orthotopic tumors at the 4th mammary fat pad (n = 7). Small size tumor volumes were selected for irradiation on day 5 (20-40 mm^3^; light blue arrow), and larger range of tumor volumes at day 24 (250-800 mm^3^; red arrow). Tumors grow moderately for the first 2 weeks and exponentially thereafter. (**B**) Tumor response curve of breast cancer orthotopic tumors of large volume range irradiated with 30 Gy single fraction of either FLASH or CONV dose rates (n = 6 per group). Tumor volumes were suppressed for the first 2 weeks and regressed thereafter. There was no significant difference between FLASH and CONV in tumor growth delay of large tumor volumes (250– 800 mm³) with a single fraction of 30 Gy.

## 4. DISCUSSION

Radiotherapy is an important treatment modality for many patients with breast cancer (*20*). The role of radiotherapy in breast cancer care is so broad that there are projected to be 2.01 million breast cancer survivors treated by radiotherapy by 2030 (*49*). Further efforts to reduce toxicity associated with radiotherapy while maintaining effectiveness would represent a significant medical advance. (*50*,*51*) Early clinical studies have been initiated. FAST-01, a feasibility study applying FLASH to bone metastases in ten patients with short term follow-up has been completed (*52*). Other studies in humans are underway, such as: FAST-02, a trial aimed at exploring thoracic bone metastases (*53*); a Phase I trial determining the maximum tolerated dose of FLASH for melanoma skin metastases (*54*); and, a randomized Phase II trial comparing FLASH and conventional radiotherapy for treating basal and squamous cell skin cancers (*55*,*56*). At the time this manuscript was written, there were no human trials identified in clinicaltrials.gov applying FLASH to primary human breast cancer.

We developed this dose-finding study to assess the ability of FLASH to eradicate small breast tumor nodules as might be seen after breast conserving surgery as well as reconfirm previously published preclinical studies using FLASH to delay tumor growth in larger, murine tumors. Our study model included Py117 cells, a breast cancer cell line syngeneic with C57BL/6 mice. Most importantly, our results demonstrated equivalent tumor control with single fraction doses of FLASH *vs* CONV at all doses utilized and with all tumor sizes.

Tumor eradication was best achieved in our small tumor cohorts at 30 Gy, the highest dose tested, and when a wider treatment margin was applied around the tumor bed. Higher single fraction doses, different orthotopic locations (such as 3^rd^ mammary fat pad), heterotopic location, and/or multiple fractionated treatment using FLASH could provide tumor eradication without the skin toxicity we noted in the 30 Gy cohort. Interestingly, while we noted skin toxicity with 30 Gy FLASH in this study, our group previously did not see lethal skin toxicity at 30 Gy when radiotherapy was applied to the hind limb (*19*). We also failed to see skin toxicity in a model of unilateral, left chest wall irradiation with 6 months of follow-up (unpublished). For this study, we elected to use the 4^th^ mammary fat pad in order assess the anti-tumor effect of FLASH *vs* CONV with larger tumors in an orthotopic location: it would not have been feasible to grow larger tumors if we had used the 3^rd^ mammary fat pad as this would have impaired the mouses’ motility and led to euthanasia of study animals before our tests could be completed. While it would seem that irradiation of the left 4^th^ mammary fat pad might have led to loss of study animals due to gastrointestinal (GI) toxicity, we did not observe signs of such toxicity.

In working with smaller tumors, we found that assessment of nodules below 50 mm^3^ potentially misleading due to misinterpretation of enlarged lymph nodes as tumor recurrence in a few study animals, confirmed on pathology. In the future, fine needle aspiration, ultrasound, or longer follow up periods will help determine the etiology of small tumor nodules without the need to euthanize study animals.

The role of treatment tumor margins (i.e., planning treatment margin) underlines the importance of beam geometry in dose delivery. The initial larger margin used with different beam geometries for FLASH and CONV highlighted the necessity of precise dosimetry. This is particularly important at the field edge where dose fall-off is critical for sparing normal tissue while ensuring adequate tumor coverage. The design of the experimental geometry in our study was necessary to address the potential for GI toxicity. By orienting the FLASH beam from floor to ceiling and positioning the mice laterally, we strategically targeted the tumor-bearing fourth mammary fat pad. This approach ensured that the tumors were within the radiation field while minimizing the abdominal exposure to radiation. While reducing the tumor irradiation margin could reduce the skin toxicity and gastrointestinal exposure, it increased the possibility that tumors might not be irradiated in their entirety, which likely contributed to more recurrences in the cohort of small tumors receiving 30 Gy treated with the 5 mm margin. This echoes the delicate balance that must be struck in radiation therapy between tumor eradication and the preservation of quality of life through the mitigation of treatment-related side effects.

The absence of statistical significance for anti-tumor effectiveness within small and large tumor cohorts between FLASH and CONV remained even after accounting for histological corrections, and the exclusion of palpable lymph nodes from tumor volume measurements. Tumor growth delay between FLASH and CONV for large size tumors was comparable, validating previous findings (*16*).

Quantitative skin toxicity experiments were not performed in this study because we have previously demonstrated skin radiotoxicity benefits of FLASH versus CONV radiation in the area of the thoracic cavity (*19*). Considering that in the current study we have irradiated tumors adjacent to the abdominal cavity we have encountered losses in mice possibly due to the radiotoxicity of the GI track, which cannot be confirmed due to the lack of postmortem biopsies. Currently, the FLASH configurations and geometries are limited to single beam angle, sustaining relevant dose rates. Ideas of different configurations like FLASH EXACT (*57*,*58*) that would introduce conformal FLASH radiotherapy would allow higher doses to the tumor without the limitation of the toxicity and even higher doses could be tested on mice without concerns of radiotoxicity. Stereotactic regimens using multiple tangents may also produce tumor control with less toxicity (*59*). A recent pilot study comparing single fraction *vs* hypofractionated radiotherapy did not, however, demonstrate improved tumor control or less toxicity, though a dose of only 20 Gy was used and tumor growth delay was the endpoint rather than tumor eradication (*41*).

The loss of a small number of mice during the study, aside from those clearly related to skin toxicity, remains unexplained. In future studies, all mice will undergo autopsies

Different tumor cell lines in immunocompetent mice as well as xenografts from humans may reveal different results based on dose and treatment margins. This highlights the need for further assessment of breast cancer eradication using larger cohorts in different tumor models to better inform the potential use of FLASH in the adjuvant therapy of human breast cancer. Further studies of both acute and late FLASH toxicity are also warranted. In addition, the radiobiology of FLASH *vs* CONV in breast cancer, as well as other solid tumors, remains incompletely understood (*18*,*60*,*61*). We are planning further in-vivo studies to understand FLASH *vs* CONV more fully given the evolving understanding of how radiotherapy interacts with the immune system and perhaps other cells and pathways associated with tumor cell death in response to therapeutic radiotherapy (*41*,*45*,*62*).

## 5. CONCLUSIONS

Single fraction 30 Gy FLASH and CONV were equally able to eradicate small tumor nodules, as might be seen surrounding a typical lumpectomy cavity, in a model of murine breast cancer. Future single-fraction studies with different cell lines, tumors, and mouse strains, both immunocompetent and immunodeficient will further inform efforts to assess the merits of FLASH *vs* CONV in human breast cancer.

## AUTHOR FOR EDITORIAL CORRESPONDENCE

Stavros Melemenidis, D.Phil.

## AUTHOR CONTRIBUTIONS

All authors met the International Committee of Medical Journal Editors (ICMJE) criteria for authorship (https://www.icmje.org/recommendations/browse/roles-and-responsibilities/defining-the-role-of-authors-and-contributors.html). Stavros Melemenidis is the first author, and Frederick M. Dirbas is senior/corresponding author.

## ABBREVIATIONS

BC: Breast Cancer
CAD: Computer-Aided Design
CONV: Conventional dose rate (radiotherapy)
DCIS: Ductal Carcinoma In Situ
EBT3: A type of radiochromic film used for dosimetry
FLASH: Ultra-high dose rate (radiotherapy)
FWHM: Full Width Half Maximum
GI: Gastrointestinal
Gy: Gray (unit of radiation dose)
IACUC: Institutional Animal Care and Use Committee
MRI: Magnetic Resonance Imaging
MMTV: Mouse Mammary Tumor Virus
NIH: National Institutes of Health
NT: Normal Tissue
OD: Optical Density
PBS: Phosphate Buffered Saline
PDD: Percentage Dose Depth
PyMT: Polyoma Middle T antigen
RT: Radiotherapy
TCD_50_: Tumor Control Dose to achieve 50% tumor eradication
TNBC: Triple-Negative Breast Cancer

## Supplementary Information

**SUP FIG 1.**
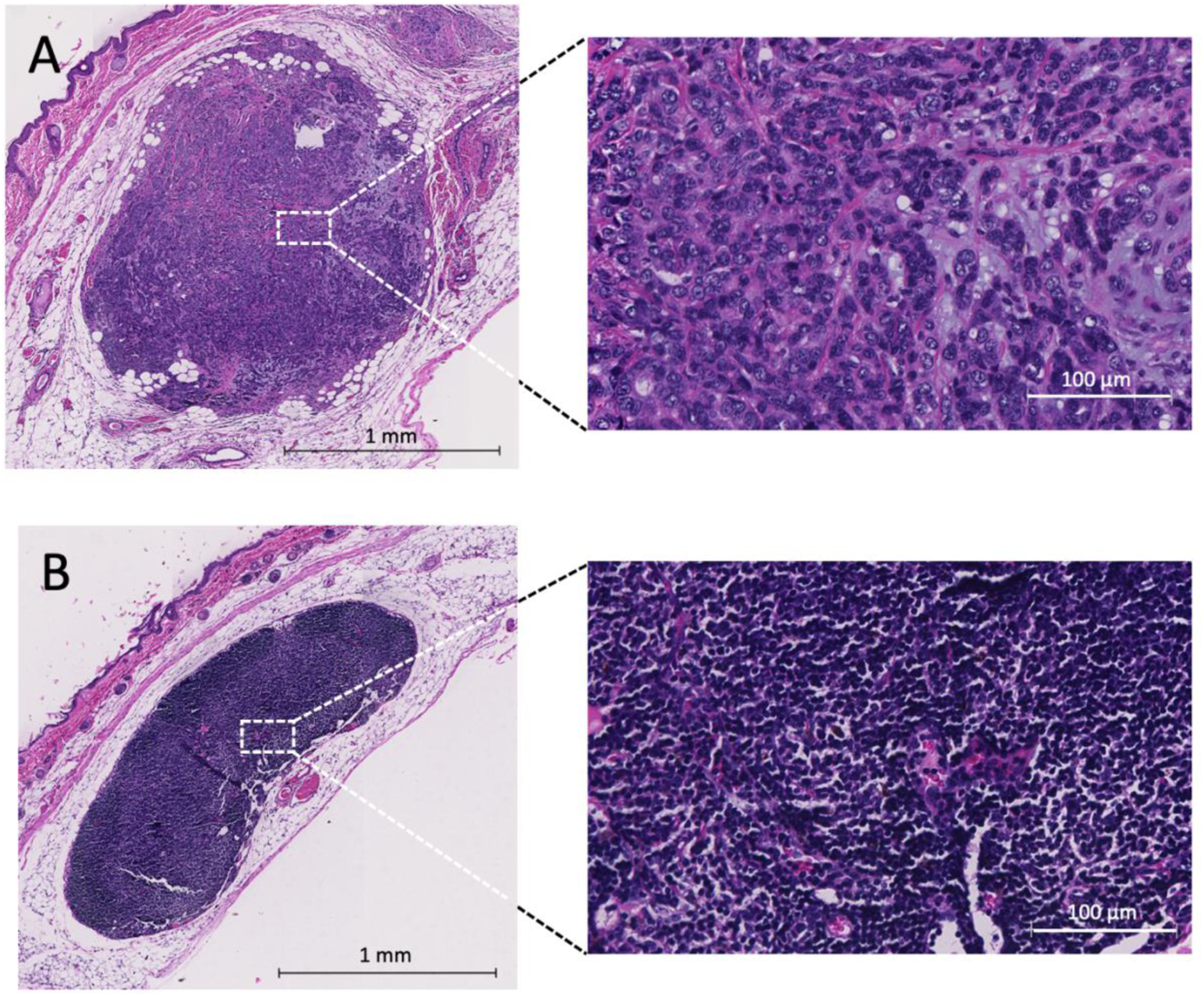
Characterization of mammary gland tissue and lymph node post-exposure to irradiation at day 46. (**A**) Histological analysis of a skin stripe from the fourth mammary fat pad, exhibiting a Py117 breast cancer orthotopic tumor. The presented photomicrograph, stained with Hematoxylin and Eosin, displays the tumor, which has a caliper-measured volume of 41.83 mm³. Blow up image highlights the cellular phenotype within the tumor. (**B**) Histological view of a lymph node situated in the implantation region of the Py117 breast cancer orthotopic tumor. Despite being accounted for as part of the tumor volume, with a caliper-measured volume of 24.63 mm³, the lymph node did not exhibit the presence of breast cancer cells upon histopathological evaluation. Blow up image highlights the cellular phenotype of the lymph node. At very small tumor volume measurements with calipers, an inflamed lymph node can be a source of false positive for residual tumor and contribute to the noise in small tumor volume measurements.

## REFERENCES

1. Fisher B. Lumpectomy and radiation therapy for the treatment of intraductal breast cancer: findings from National Surgical Adjuvant Breast and Bowel Project B-17. J Clin Oncol. 1998;16(2):441–52.

2. Dirbas FM, Jeffrey SS, Goffinet DR. The evolution of accelerated, partial breast irradiation as a potential treatment option for women with newly diagnosed breast cancer considering breast conservation. Vol. 19. 2004. p. 673–705.

3. Shah C. Outcomes with Partial Breast Irradiation vs. Whole Breast Irradiation: a Meta-Analysis. Ann Surg Oncol; 2021.

4. Barazzuol L, Coppes RP, Luijk P. Prevention and treatment of radiotherapy-induced side effects. Vol. 14. Mol Oncol; 2020. 1538–1554 p.

5. Williams PA. Patient-reported outcomes of the relative severity of side effects from cancer radiotherapy. Support Care Cancer. 2020;28(1):309–16.

6. Milam EC, Rangel LK, Pomeranz MK. Dermatologic sequelae of breast cancer: From disease, surgery, and radiation. Int J Dermatol. 2021;60(4):394–406.

7. Magill LJ. Determining the outcomes of post-mastectomy radiation therapy delivered to the definitive implant in patients undergoing one- and two-stage implant-based breast reconstruction: A systematic review and meta-analysis. J Plast Reconstr Aesthet Surg. 2017;70(10):1329–35.

8. Martin S. Autologous Fat Grafting and the Occurrence of Radiation-Induced Capsular Contracture. Ann Plast Surg. 2021;86(5S Suppl 3):414–7.

9. Chen JJ. Development of a Classification Tree to Predict Implant-Based Reconstruction Failure with or without Postmastectomy Radiation Therapy for Breast Cancer. Vol. 28. 2021. p. 1669–79.

10. Cordeiro PG. The impact of postmastectomy radiotherapy on two-stage implant breast reconstruction: an analysis of long-term surgical outcomes, aesthetic results, and satisfaction over 13 years. Vol. 134. Plast Reconstr Surg; 2014. 588–595 p.

11. Hughes KS. Lumpectomy plus tamoxifen with or without irradiation in women 70 years of age or older with early breast cancer. N Engl J Med. 2004;351(10):971–7.

12. Shah C. The Clinical Utility of DCISionRT((R)) on Radiation Therapy Decision Making in Patients with Ductal Carcinoma In Situ Following Breast-Conserving Surgery. 2021.

13. Hamidi M, Moody JS, Kozak KR. Refusal of radiation therapy and its associated impact on survival. Am J Clin Oncol. 2010;33(6):629–32.

14. Tuttle TM. Omission of radiation therapy after breast-conserving surgery in the United States: a population-based analysis of clinicopathologic factors. Cancer. 2012;118(8):2004– 13.

15. Blanpain C. DNA-damage response in tissue-specific and cancer stem cells. Cell Stem Cell. 2011;8(1):16–29.

16. Favaudon V, Caplier L, Monceau V, et al. Ultrahigh dose-rate FLASH irradiation increases the differential response between normal and tumor tissue in mice. Sci Transl Med. 2014 Jul 16;6(245):245ra93.

17. Wilson JD, Hammond EM, Higgins GS, et al. Ultra-High Dose Rate (FLASH) Radiotherapy: Silver Bullet or Fool’s Gold? Front Oncol [Internet]. 2020 Jan 17 [cited 2020 Mar 18];9. Available from: https://www.ncbi.nlm.nih.gov/pmc/articles/PMC6979639/

18. Spitz DR. An integrated physico-chemical approach for explaining the differential impact of FLASH versus conventional dose rate irradiation on cancer and normal tissue responses. Radiother Oncol; 2019. 139 23–27.

19. Soto LA, Casey KM, Wang J, et al. FLASH Irradiation Results in Reduced Severe Skin Toxicity Compared to Conventional-Dose-Rate Irradiation. Radiat Res. 2020 Dec 1;194(6):618–24.

20. DeSantis CE. Breast cancer statistics, 2019. CA Cancer J Clin. 2019;69(6):438–51.

21. Rohrer Bley C. Dose and volume limiting late toxicity of FLASH radiotherapy in cats with squamous cell carcinoma of the nasal planum and in mini-pigs. Clin Cancer Res. 2022;

22. Suo M, Shen H, Lyu M, et al. Biomimetic Nano-Cancer Stem Cell Scavenger for Inhibition of Breast Cancer Recurrence and Metastasis after FLASH-Radiotherapy. Small. 2024 Jul;20(29):e2400666.

23. Zhu H, Xie DH, Yang Y, et al. The Immune Response and Intestinal Injury after X-Ray FLASH Irradiation in Murine Breast Cancer Transplanted Models. International Journal of Radiation Oncology*Biology*Physics [Internet]. 2022 Nov 1 [cited 2024 Aug 18];114(3, Supplement):S66–7. Available from: https://www.sciencedirect.com/science/article/pii/S0360301622011786

24. Lattery G, Kaulfers T, Cheng C, et al. Pencil Beam Scanning Bragg Peak FLASH Technique for Ultra-High Dose Rate Intensity-Modulated Proton Therapy in Early-Stage Breast Cancer Treatment. Cancers (Basel) [Internet]. 2023 Sep 14 [cited 2024 Aug 18];15(18):4560. Available from: https://www.ncbi.nlm.nih.gov/pmc/articles/PMC10527307/

25. Franciosini G, Carlotti D, Cattani F, et al. IOeRT conventional and FLASH treatment planning system implementation exploiting fast GPU Monte Carlo: The case of breast cancer. Physica Medica [Internet]. 2024 May 1 [cited 2024 Aug 18];121:103346. Available from: https://www.sciencedirect.com/science/article/pii/S1120179724001418

26. van Marlen P, van de Water S, Dahele M, et al. Single Ultra-High Dose Rate Proton Transmission Beam for Whole Breast FLASH-Irradiation: Quantification of FLASH-Dose and Relation with Beam Parameters. Cancers (Basel). 2023 Apr 30;15(9):2579.

27. Adrian G. Cancer Cells Can Exhibit a Sparing FLASH Effect at Low Doses Under Normoxic In Vitro-Conditions. Vol. 11. Front Oncol; 2021. 686142 p.

28. Rosen PP, Fracchia AA, Urban JA, et al. “Residual” mammary carcinoma following simulated partial mastectomy. Cancer. 1975 Mar;35(3):739–47.

29. Lagios MD. Multicentricity of breast carcinoma demonstrated by routine correlated serial multicentric breast carcinoma subgross and radiographic examination. Cancer [Internet]. 1977 [cited 2024 May 23];40(4):1726–34. Available from: https://onlinelibrary.wiley.com/doi/abs/10.1002/1097-0142%28197710%2940%3A4%3C1726%3A%3AAID-CNCR2820400449%3E3.0.CO%3B2-O

30. Renton SC, Gazet JC, Ford HT, et al. The importance of the resection margin in conservative surgery for breast cancer. Eur J Surg Oncol. 1996 Feb;22(1):17–22.

31. Fisher B. Twenty-year follow-up of a randomized trial comparing total mastectomy, lumpectomy, and lumpectomy plus irradiation for the treatment of invasive breast cancer. N Engl J Med. 2002;347(16):1233–41.

32. Clark RM, Whelan T, Levine M, et al. Randomized clinical trial of breast irradiation following lumpectomy and axillary dissection for node-negative breast cancer: an update. Ontario Clinical Oncology Group. J Natl Cancer Inst. 1996 Nov 20;88(22):1659–64.

33. Forrest AP, Stewart HJ, Everington D, et al. Randomised controlled trial of conservation therapy for breast cancer: 6-year analysis of the Scottish trial. Scottish Cancer Trials Breast Group. Lancet. 1996 Sep 14;348(9029):708–13.

34. Liljegren G, Holmberg L, Bergh J, et al. 10-Year results after sector resection with or without postoperative radiotherapy for stage I breast cancer: a randomized trial. J Clin Oncol. 1999 Aug;17(8):2326–33.

35. Holli K, Saaristo R, Isola J, et al. Lumpectomy with or without postoperative radiotherapy for breast cancer with favourable prognostic features: results of a randomized study. Br J Cancer. 2001 Jan;84(2):164–9.

36. Veronesi U, Orecchia R, Maisonneuve P, et al. Intraoperative radiotherapy versus external radiotherapy for early breast cancer (ELIOT): a randomised controlled equivalence trial. Lancet Oncol. 2013 Dec;14(13):1269–77.

37. Vicini FA, Cecchini RS, White JR, et al. Long-term primary results of accelerated partial breast irradiation after breast-conserving surgery for early-stage breast cancer: a randomised, phase 3, equivalence trial. Lancet. 2019 Dec 14;394(10215):2155–64.

38. Chen HL, Zhou JQ, Chen Q, et al. Comparison of the sensitivity of mammography, ultrasound, magnetic resonance imaging and combinations of these imaging modalities for the detection of small (≤2 cm) breast cancer. Medicine (Baltimore). 2021 Jul 2;100(26):e26531.

39. Langer SA, Horst KC, Ikeda DM, et al. Pathologic correlates of false positive breast magnetic resonance imaging findings: which lesions warrant biopsy? The American Journal of Surgery [Internet]. 2005 Oct 1 [cited 2024 Jun 21];190(4):633–40. Available from: https://www.sciencedirect.com/science/article/pii/S0002961005005647

40. Sørensen BS, Sitarz MK, Ankjærgaard C, et al. Pencil beam scanning proton FLASH maintains tumor control while normal tissue damage is reduced in a mouse model. Radiotherapy and Oncology [Internet]. 2022 Oct [cited 2023 Aug 24];175:178–84. Available from: https://linkinghub.elsevier.com/retrieve/pii/S0167814022002547

41. Almeida A, Godfroid C, Leavitt RJ, et al. Antitumor Effect by Either FLASH or Conventional Dose Rate Irradiation Involves Equivalent Immune Responses. Int J Radiat Oncol Biol Phys [Internet]. 2024 Mar 15 [cited 2024 Jun 21];118(4):1110–22. Available from: https://www.ncbi.nlm.nih.gov/pmc/articles/PMC11093276/

42. Aguilera TA. Reprogramming the immunological microenvironment through radiation and targeting Axl. Nat Commun. 2016;7:13898.

43. Wang J, Melemenidis S, Manjappa R, et al. Dosimetric calibration of an anatomically specific ultra-high dose rate electron irradiation platform for preclinical FLASH radiobiology experiments [Internet]. arXiv; 2023 [cited 2024 May 23]. Available from: http://arxiv.org/abs/2312.10632

44. Schuler E. Experimental Platform for Ultra-high Dose Rate FLASH Irradiation of Small Animals Using a Clinical Linear Accelerator. Int J Radiat Oncol Biol Phys. 2017;97(1):195– 203.

45. Levy K. Abdominal FLASH irradiation reduces radiation-induced gastrointestinal toxicity for the treatment of ovarian cancer in mice. Sci Rep. 2020;10(1):21600.

46. Eggold JT, Chow S, Melemenidis S, et al. Abdominopelvic FLASH Irradiation Improves PD-1 Immune Checkpoint Inhibition in Preclinical Models of Ovarian Cancer. Mol Cancer Ther. 2022 Feb;21(2):371–81.

47. Drayson OG, Melemenidis S, Katila N, et al. A multi-institutional study to investigate the sparing effect after whole brain electron FLASH in mice: Reproducibility and temporal evolution of functional, electrophysiological, and neurogenic endpoints [Internet]. bioRxiv; 2024 [cited 2024 Apr 8]. p. 2024.01.25.577164. Available from: https://www.biorxiv.org/content/10.1101/2024.01.25.577164v1

48. Lewis D, Micke A, Yu X, et al. An efficient protocol for radiochromic film dosimetry combining calibration and measurement in a single scan. Med Phys. 2012 Oct;39(10):6339– 50.

49. Bryant AK. Trends in Radiation Therapy among Cancer Survivors in the United States, 2000-2030. Vol. 26. Cancer Epidemiol Biomarkers Prev; 2017. 963–970 p.

50. Wu Y (Fred), No HJ, Breitkreutz DY, et al. Technological Basis for Clinical Trials in FLASH Radiation Therapy: A Review. ARO [Internet]. 2021 Jun 1 [cited 2023 Jul 14];6–14. Available from: https://www.crossref.org/

51. Loo BW, Verginadis II, Sørensen BS, et al. Navigating the Critical Translational Questions for Implementing FLASH in the Clinic. Seminars in Radiation Oncology [Internet]. 2024 Jul 1 [cited 2024 Jun 21];34(3):351–64. Available from: https://www.sciencedirect.com/science/article/pii/S1053429624000286

52. Mascia AE, Daugherty EC, Zhang Y, et al. Proton FLASH Radiotherapy for the Treatment of Symptomatic Bone Metastases: The FAST-01 Nonrandomized Trial. JAMA Oncology [Internet]. 2023 Jan 1 [cited 2024 Jun 21];9(1):62–9. Available from: 10.1001/jamaoncol.2022.5843

53. Study Details | FLASH Radiotherapy for the Treatment of Symptomatic Bone Metastases in the Thorax | ClinicalTrials.gov [Internet]. [cited 2024 Aug 18]. Available from: https://clinicaltrials.gov/study/NCT05524064

54. Study Details | Irradiation of Melanoma in a Pulse | ClinicalTrials.gov [Internet]. [cited 2024 Aug 18]. Available from: https://clinicaltrials.gov/study/NCT04986696

55. Kinj R, Gaide O, Jeanneret-Sozzi W, et al. Randomized phase II selection trial of FLASH and conventional radiotherapy for patients with localized cutaneous squamous cell carcinoma or basal cell carcinoma: A study protocol. ctRO [Internet]. 2024 Mar 1 [cited 2024 Jun 21];45. Available from: https://www.ctro.science/article/S2405-6308(24)00020-X/fulltext

56. Study Details | FLASH Radiotherapy for Skin Cancer | ClinicalTrials.gov [Internet]. [cited 2024 Aug 18]. Available from: https://clinicaltrials.gov/study/NCT05724875

57. Ko RB, Soto LA, von Eyben R, et al. Evaluating the Reproducibility of Mouse Anatomy under Rotation in a Custom Immobilization Device for Conformal FLASH Radiotherapy. Radiat Res [Internet]. 2020 Dec 1 [cited 2023 Nov 1];194(6):600–6. Available from: https://www.ncbi.nlm.nih.gov/pmc/articles/PMC7856226/

58. Maxim PG, Tantawi SG, Loo BW. PHASER: A platform for clinical translation of FLASH cancer radiotherapy. Radiother Oncol. 2019 Oct;139:28–33.

59. Fisher B, Dignam J, Wolmark N, et al. Lumpectomy and radiation therapy for the treatment of intraductal breast cancer: findings from National Surgical Adjuvant Breast and Bowel Project B-17. J Clin Oncol. 1998 Feb;16(2):441–52.

60. Friedl AA. Radiobiology of the FLASH effect. Med Phys. 2022;49(3):1993–2013.

61. Buchsbaum JC. FLASH Radiation Therapy: New Technology Plus Biology Required. Int J Radiat Oncol Biol Phys. 2021;110(4):1248–9.

62. Rama N. Improved Tumor Control Through T-cell Infiltration Modulated by Ultra-High Dose Rate Proton FLASH Using a Clinical Pencil Beam Scanning Proton System. 2019.

